# Optimization of modularity during development to simplify walking control across strides

**DOI:** 10.1101/2022.11.07.515149

**Authors:** Elodie Hinnekens, Bastien Berret, Estelle Morard, Manh-Cuong Do, Marianne Barbu-Roth, Caroline Teulier

## Abstract

Walking in adults seems to rely on a small number of modules allowing to reduce the number of degrees of freedom effectively regulated by the central nervous system (CNS). However, the extent to which modularity evolves during development remains unknown, particularly regarding the ability to generate several strides in an optimized manner. Here we compared the modular organization of toddlers and adults during several strides of walking. We recorded the electromyographic activity of 10 bilateral (lower limbs) muscles in adults (n=12) and toddlers (n=12) during 8 gait cycles, and used non-negative matrix factorization to model the underlying modular command. While the muscular activity of all strides could be factorized into a consistent low-dimensional modular organization in adults, significantly more computational modules were needed in toddlers to account for their greater stride-by-stride variability. Activations of these modules varied more across strides and was less parsimonious in toddlers than in adults, even when balances constrained were diminished. These findings suggest that the modular control of locomotion of adults evolves as the organism develops and practices. They also suggest that new walker can flexibly activate a higher number of modules and benefit from a higher space of possible action, which could serve motor exploration.

## INTRODUCTION

Walking is easily performed by adult organisms despite the abundance of degrees of freedoms (DOFs) that the central nervous system (CNS) has to deal with (Bernstein, 1967). This antagonism has led to propose that such complex motor behaviors could be generated by the activation of a small number of “modules” stored in the CNS (Bizzi et al., 1991; Mussa-Ivaldi et al., 1994; d’Avella et al., 2003). A module, also called building-block, motor primitive or muscle synergy in the literature, is a neural structure in charge of producing a specific muscle pattern when activated by higher centers (Bizzi et al., 2008). Modularity would thus simplify the organization of behavior by reducing the effective number of DOFs controlled by the CNS (Bizzi and Cheung, 2013). Studying walking in human adults, researchers identified that the electromyographic (EMG) activity of walking could be factorized into a few spatial and temporal computational modules that are suspected to correspond to actual physiological modules and to simplify the motor command of walking (Ivanenko et al., 2004; Chvatal and Ting, 2013).

This simplified modular control is concomitant with an optimized and adaptable gait (Alexander, 1991; Choi and Bastian, 2007; Neptune et al., 2009; Chvatal and Ting, 2012). However, the ease with which we move has developed over years and years of practice. Walking is indeed more difficult to handle in new walkers who fall on average 17 times per hour (Adolph et al., 2013) and who do not yet owe mature gait patterns (Sutherland, 1997; Lacquaniti et al., 2012a). Particularly, walking patterns of toddlers still differ from that of adults because of the high variability of muscle activity across steps (Chang et al., 2006; Teulier et al., 2012). A growing body of evidence suggests that such variability is centrally and purposively generated in novices and of great interest for motor learning as it allows motor exploration (Kao et al., 2008; Mandelblat-Cerf et al., 2009; Ziegler et al., 2010; Wu et al., 2014; Dhawale et al., 2017). Variability could also be generated by feedback corrections to control body weight (Kerkman et al., 2022). Yet, toddlers’ motor patterns remain instable and immature even with balance support, thereby suggesting that stride-by-stride variability does not specifically originate from balance-related processes (Ivanenko et al., 2005).

Inferring modules from averaged or single-step data, researchers found that the number of modules increased from birth to toddlerhood and then stabilized until adulthood (Dominici et al., 2011; Sylos-labini et al., 2020). However, whether several steps of toddlers walking could also be factorized into a simplified modular control as adults remain unknown. Although the space of motor solutions in inherently restricted by a low-dimensional system (Valero-Cuevas, 2009; Cohn et al., 2018), a substantial trial-by-trial variability of the muscular command could still be generated from a few modules if the modules are themselves activated erratically (Hinnekens et al., 2022). To clarify this issue, we thoroughly investigated the intricate link between stride-by-stride variability of muscle activity and muscle modularity in toddlers and adults. In particular, we characterized the computational modules that account for muscle patterns across several strides in both populations and assessed the consequences of data averaging and of balance control. As learning in a modular system consists both in learning modules and controlling their parameters (d’Avella and Pai, 2010), we systematically analyzed the dimensionality of the inferred modular systems (i.e. number of modules) as well as the properties of their activations parameters (variability and selectivity of modules’ activations).

## METHODS

### Experimental Protocol

Twelve adults (7 females, 5 males, age 25.8 ± 4 years [mean ± SD]) and twelve toddlers were recruited for this study (3 females, 9 males, age 15,5 ± 2 months). Toddler walking was recorded at a maternity while adults were tested in a laboratory of the university. Toddler experiment was planned one to five weeks after parents reported they ability to walk independently and unsupported as a main mode of locomotion. 11 parents were able to give us the exact day when their child was able to “cross an entire room of about sixteen feet by walking”. Hence, toddlers walking experience when coming to the lab was 19.3 ± 7.1 days (mean ± SD). The protocol was in accordance with the Helsinki Declaration and approved by the French Committee of People Protection. Parents of the children as well as adults’ participants gave written and informed consent before participation. Adults and toddlers were asked to walk barefoot at a comfortable speed during approximately one minute. Toddlers had to walk back and forth along a two meters exercise mat following a linear trajectory without any help from adults. About 8 steps could be recorded for a single straight walking path. Breaks were taken by toddlers when needed. In order to compare walking in toddlers with a control condition that would involve less balance constraints, we also held 10 of the toddlers while stepping on a treadmill (see methods – control conditions).

### Data recording

#### EMG recording

Ten bilateral muscles were recorded for this study, as previous investigations of toddlers modularity identified bilateral modules (Dominici et al., 2011; Sylos-labini et al., 2020). We recorded the activity of muscles from shanks thighs and buttocks: tibialis anterior, soleus, rectus femoris, biceps femoris and gluteus medius. Another 6 extra muscles (involving trunk and proximal upper limb regions) were recorded for the need of other studies. Electrodes were placed according to SENIAM recommendations (Surface EMG for Non-Invasive Assessment of Muscles, seniam.org). Surface EMG data were recorded with the Cometa system (Biometrics ®) at 2000 Hz with bipolar electrodes (21×41 millimeters).

#### Motion capture

Motion was recorded in adults and toddlers in order to detect stride events. In adults, we used an eight-camera Qualysis® system recording at 100 Hz. Nineteen markers were placed on each individuals, of which we used heel and toe (second metatarsal head) ones to determine gait events (as in O’Connor et al. 2007). The motion capture system was synchronized with the EMG systems thanks to a common trigger. In toddlers, we used two 2D cameras recording at 50 Hz. The same trigger as in adult recording was used to launch these cameras and the EMG systems. The toddler had to go back and forth along a two meters exercise mat; thus, cameras were placed on each side of the mat so we would have a clear vision of both sides of the body to detect gait events.

### Data processing and computed parameters

#### Identification of stride events

We identified the Foot Off and Foot Strike events in both populations. A stride was defined from a Foot Off event to another Foot Off event. In both populations, we considered only right, alternated strides. Strides were analyzed only when the toddler was following a linear trajectory. The first and last steps of a crossing were never considered.

In adults we used the foot velocity algorithm described by O’Connor et al. (2007) to detect Foot Off and Foot Strike. These events are more often referred to as Toe Off and Heel Strike in adults but we call them Foot Off and Foot Strike because toddlers do not necessarily start swing with the toe or end swing with the heel.

In toddlers, a trained coder screened all the videos and identified the Foot Off and Foot Strike events, as done in Teulier et al. (2012). The Foot Off event was defined as the last frame before the whole foot would stop touching the floor. The Foot Strike event was defined as the first frame where any part of the foot would touch the floor. The same coder waited a month after having done these identifications and identified again fifty strides from five different toddlers in order to compute an Intraclass Correlation Coefficient (ICC). This ICC was 0.99 showing excellent reliability in gait events identification. Figure 1 shows the view from one camera during a whole stride.

**Figure 1.**
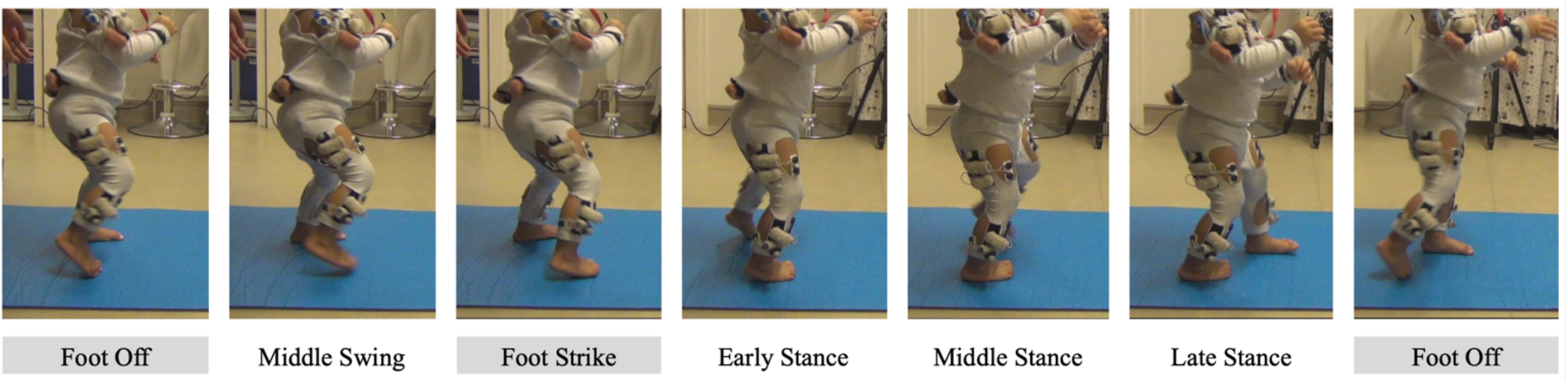
Illustration of stride events taken by the toddler. Events identified by the coder are shown with a grey background.

As our analysis focuses on intra-individual variability, it was important to analyze the same number of strides in each participant. For some toddlers getting a lot of straight steps were hard, hence 8 strides were the number of strides we could get for each participant. From those, we computed three basic kinematic parameters: stride duration, standard deviation of stride duration and proportion of swing and stance phases.

#### EMG processing

Filtering and interpolation of data was done as reported by Ivanenko et al. (2013). Data were high-passed filtered (40 Hz, zero lag fourth order Butterworth filter), rectified, and low-pass filtered (10 Hz, zero lag fourth order Butterworth filter). For each gait cycle, the signal was interpolated to 200 time points. The signal of each muscle was normalized by its maximum amplitude across 8 strides. In each participant, we obtained a *(t × s) × m* matrix where *t* equaled 200 time points, *s* equaled 8 strides and *m* equaled 10 muscles. Each entry took values between 0 and 1, 1 representing the normalized maximum activity of the corresponding muscle. These matrices constituted the non-averaged EMG signals. Then we created other matrices from the latter ones by averaging across strides. These matrices were of the form *t × m* and constituted the averaged EMG signals. As no consensus exists regarding temporal normalization of data that precedes non-negative matrix factorization, we verified the results of the study after having computed different variants (phase-interpolation rather than cycle-interpolation, computing RMS rather than standard interpolation, or basing the interpolation on 20 time-points rather than 200).

From the non-averaged EMG signals, we computed an index of EMG variability, as was done in Hinnekens et al. (2020). This index was defined as the standard deviation computed point by point from the pre-processed EMG and across the 8 strides.

As cross-talk might be an issue when recording surface EMG data, we used the same criterion than Dominici et al. (2011) to report potential cross-talk (Pearson correlation coefficient >0.2). We computed Pearson correlation coefficients across four pairs of muscles on each side (rectus femoris and biceps femoris, tibialis anterior and soleus, gluteus medius and rectus femoris, gluteus medius and biceps femoris) on data after having applied the high-pass filter only. Thus, this analysis was made for 8 pairs of muscles, 8 strides and 12 subjects in both populations (i.e. 1536 samples). 6% of samples had a correlation coefficient >0.2. For these samples, we checked whole recordings and verified that different strides from one recording did not have the same correlation coefficient and were not all >0.2.

#### EMG Factorization

We used the Space-by-Time Decomposition method in order to factorize the signal into spatial and temporal modules. This method, unifying previous approaches as described in Delis et al. (2014), is based on non-negative matrix factorization (NNMF). It allows to extract both spatial and temporal EMG invariants (modules) while retaining intra-individual variability in a low dimensional space (activation coefficients). Therefore, it allows to directly test the hypothesis that toddlers would benefit from the same low dimensional modular organization than adults while producing variability by generating differences within activation parameters of modules. Precisely, the EMG activity is factorized so that any muscle pattern ***m***_***s***_(*t*) of the stride s is considered as the following double linear combination of invariant spatial and temporal modules:

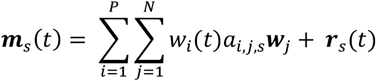

where *P* and *N* are the numbers of temporal and spatial modules respectively, ***w***_*i*_(*t*) and ***w***_*j*_ are the temporal and spatial modules respectively, *a*_*i,j,s*_ is a scalar activation coefficient (function of the pair of modules it activates and stride *s*), and ***r***_*s*_(*t*) is the residual reconstruction error describing the difference between the original signal and the reconstructed one. In this method, a spatial module is defined as an invariant ensemble of muscles which are activated together with different proportions (a constant 10-D vector here), and a temporal module is defined as a waveform that describes the amplitude changes of any spatial module over a gait cycle, invariant in regards to the different strides but time-varying within one stride (a time-varying function represented by 200 points here). An activation coefficient is attributed at each stride to each possible pair of spatial and temporal modules and quantifies their concurrent activation: a low activation coefficient means that the corresponding spatial and temporal modules are not activated together while a high one reveals a concurrent activation. Scalar activation coefficients are free to vary for each stride when using non-averaged EMG data. The algorithm was run with a custom Matlab® code. It starts from random guesses of the solution (modules and activation coefficients) and modifies these quantities until the reconstruction error is minimal, using a convergence criterion. This process was repeated fifty times for each decomposition to minimize the probability of being stuck in a local minimum.

#### Code accessibility

The custom code underlying the computational analysis is available at: http://hebergement.universite-paris-saclay.fr/berret/software/sNM3F.zip. See Delis et al. (2014) for details about the code and algorithm.

#### Quality of reconstruction criteria

The quality of reconstruction criteria, called Variance Accounted For (VAF), was computed as the coefficient of determination between the initial matrix of data and the reconstructed one using the following formula:

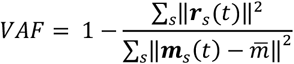

where 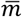 is the mean level of muscle activity across all samples and ‖·‖ represents the Frobenius norm.

The VAF quantifies how faithful the original pattern is described by the decomposition. As the purpose of the decomposition is to reduce dimensionality, the VAF is directly related to the number of modules that is extracted (the VAF increases when the number of modules increases). Hence the signals can be considered as resulting from a low-dimensional modular organization if they can be factorized into a small number of modules with a good-enough VAF, as this is the case in adults. Numerous studies indeed identified that extracting four spatial and temporal modules from walking in adults was sufficient to describe its EMG activity and resulted in biomechanically functional modules (Neptune et al., 2009; Clark et al., 2010; Lacquaniti et al., 2012b; Hinnekens et al., 2020). Therefore, we used here two complementary approaches to study the command of toddlers in comparison to the one of adults: by comparing the VAF resulting from the extraction of the same number of modules (variant I) and by comparing the necessary number of modules to get the same VAF value (variant II).

#### Variant I: extracting the same number of modules than in adults and comparing the resulting VAF

This approach directly tests the hypothesis that a low-dimensional modular command model could equally fit each dataset. Based on previous literature we considered 4 spatial and temporal modules to be an efficient low-dimensional modular command (Neptune et al., 2009; Clark et al., 2010; Lacquaniti et al., 2012b; Hinnekens et al., 2020). Thus, we extracted 4 modules from each dataset and compared the resulting VAF. Interestingly, this variant allows to quantify the extent to which a reduction of dimensionality can fit the data with a continuous variable (rather than only analyzing the discrete number of modules), which yields more precise data for the statistical analysis. This approach was applied to both averaged and non-averaged data in order to quantify the effects of intra-individual variability on modularity.

#### Variant II: computing the necessary number of modules to reach a threshold VAF and comparing features of modular organization

After having tested if a low-dimensional modularity hypothesis would fit toddlers’ data with variant I, this variant allowed to determine which dimensionality should be effectively considered in toddlers to allow a sufficient goodness of fit. We only analyzed non-averaged data from this point forward. This approach has been more commonly used in previous studies, although no consensus exists about the threshold VAF for a good quality of reconstruction (Alessandro et al., 2013). Hence here again we based the analysis on the numerous studies that identified four spatial and temporal modules as sufficient to describe the EMG activity of adult walking (Neptune et al., 2009; Clark et al., 2010; Lacquaniti et al., 2012b; Hinnekens et al., 2020) and we defined the threshold for a good quality of reconstruction as the averaged VAF (across individuals) obtained after extracting four modules from non-averaged data of adults walking (as used in Hinnekens et al. 2020 in order to compare behaviors). The resulting threshold was 0.75. Then we extracted enough modules in each toddler’s dataset to reach this threshold VAF. This led to the identification of a specific modular organization in each toddler from which we could study the features of modules activations. For this purpose, we defined two indexes to compare the characteristics of modules activations of both populations, namely the Index of Recruitment Variability (IRV) and the Index of Recruitment Selectivity (IRS).

The IRV is defined as the average standard deviations of activation coefficients across strides (the standard deviation of activation coefficients is computed for each possible pair of spatial and temporal modules, and we consider the averaged value across all possible pairs). It indicates how variable are the stride-dependent activations of pairs of temporal/spatial modules. When the IRV is lower, modules recruitment can be considered as more stable (Figure 2A), while it increases when modules recruitment is more variable across strides (Figure 2B).

**Figure 2.**
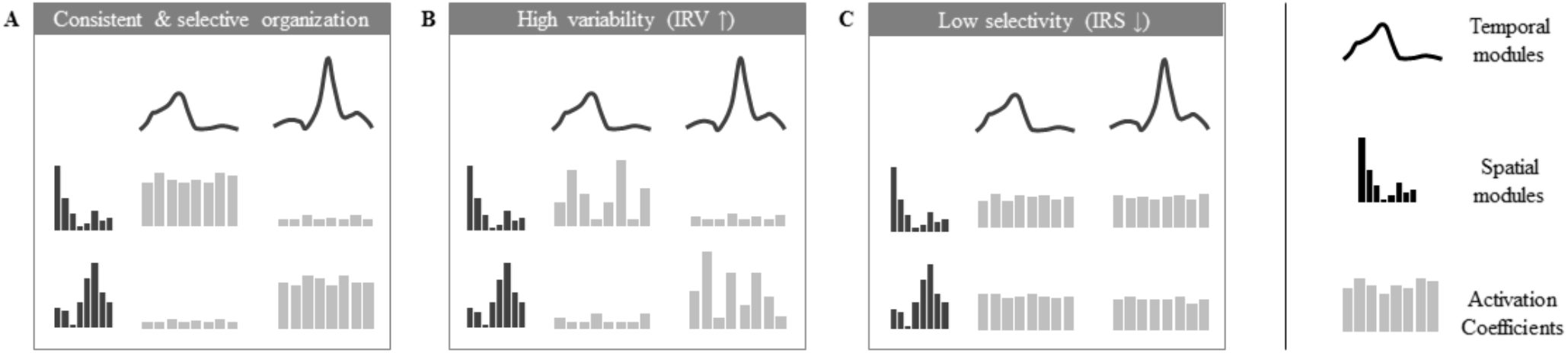
Illustration of the analyzed properties of the modular organization. Depicted are modular organizations of two spatial and temporal modules. At the center of each figure 8 activation coefficients are depicted (one for each stride). The variation of these coefficients defines the properties of consistency and selectivity. A: example of a consistent and selective modular organization (activation coefficients show low variability and are significantly higher for given pairs of spatial and temporal module). B: example of a modular organization with a high IRV (activation coefficients show high variability across strides). C: example of a modular organization with a low IRS (activation coefficients are equivalent for each possible pair of spatial and temporal modules).

The IRS is computed using a metric to assess the sparseness of activation coefficients (Hoyer, 2004). It quantifies the extent to which modules are selectively and parsimoniously recruited. It was already established that modularity of mature walking was associated with the activation of only four specific pairs of spatial and temporal modules (Hinnekens et al., 2020) where each spatial module is associated with a single temporal module (as in the example of Figure 2A with two pairs of spatial and temporal modules, which would correspond to a high IRS). In contrast a lower IRS would correspond to a non-selective command exploiting the possible multiplexing of spatial and temporal modules and resulting in the concurrent activation of more pairs of modules (Figure 2C).

#### Supplementary analyses

Following those computations, we tested the robustness of our findings regarding several methodological choices, as different methods of EMG preprocessing and EMG factorization exist within the literature. Therefore we repeated the analysis by extracting spatial modules only (Chvatal and Ting, 2013), temporal modules only (Dominici et al., 2011), and from different EMG preprocessing (band-pass filtering between 30 and 400 hz, time interpolation computed with root mean squares, and phase interpolation instead of cycle interpolation, see Hinnekens et al. 2020).

### Statistical analyses

#### Basic kinematic and EMG parameters

We compared basic kinematic and EMG parameters between the two populations: stride duration, standard deviation of stride duration, proportion of phases and index of EMG variability (described above). We used student t-test on independent samples.

#### Comparison of the goodness of fit with fixed number of modules (variant I)

To compare the faithfulness of modeling with fixed number of modules we computed the VAF resulting from the extraction of four spatial and temporal modules in both populations. To quantify the effect of stride-to-stride variability on this modeling and how it interacts with age, the extraction was made both from averaged and non-averaged signals. As VAF is not normally distributed, we transformed values using Fisher z-transformation before this statistical analysis. We compared the resulting transformed VAF with a mixed ANOVA, with one within-subjects factor (averaging or not) and one between-subjects factor (toddlers/adults). Student-t tests were performed as post-hocs and resulting p-values were multiplied by 4 to account for multiple comparisons (4 post-hoc tests).

#### Comparison of the modular parameters with fixed goodness of fit (variant II)

To identify faithful modeling of data in toddlers, we incremented the number of modules until the threshold VAF would be reach, as described above. We obtained a specific modular organization with a specific number of modules in each toddler. IRV and IRS were computed from these specific modular organizations. Values for adults and toddlers were compared thanks to a student t-test on independent samples.

To facilitate comparisons between adults and toddlers we reproduced these analyses for the modeling of toddler’s data with four spatial and temporal modules. As adult’s data are modeled with four spatial and temporal modules, this ensures that differences between modular parameters are not due to the difference in the number of modules.

### Control conditions

We used two control conditions to verify that our results were not mainly due to non-physiological variability or to variable feedback regulations that would be associated with new balance constraints in toddlers.

The first control condition called “computational control” verifies the results of adults’ data. It was created from adult walking data in order to control the fact that gait events might be less easily recognizable in toddlers. As we coded event in toddler visually, and even if the intraclass reliability was excellent, we could expect that this coding could result in less precision than the adult algorithm. Hence to check the possible effects of an unintended shifting, we introduced random offset delay in the adult event detection from -2 to +2 frames of the real time event and repeated the analysis from adult data that were cut-off according to this randomly shifted-detection matrix instead of the original one (Figure 3A). Individual data for each index in both primary and control conditions are reported in Table *3*.

**Figure 3.**
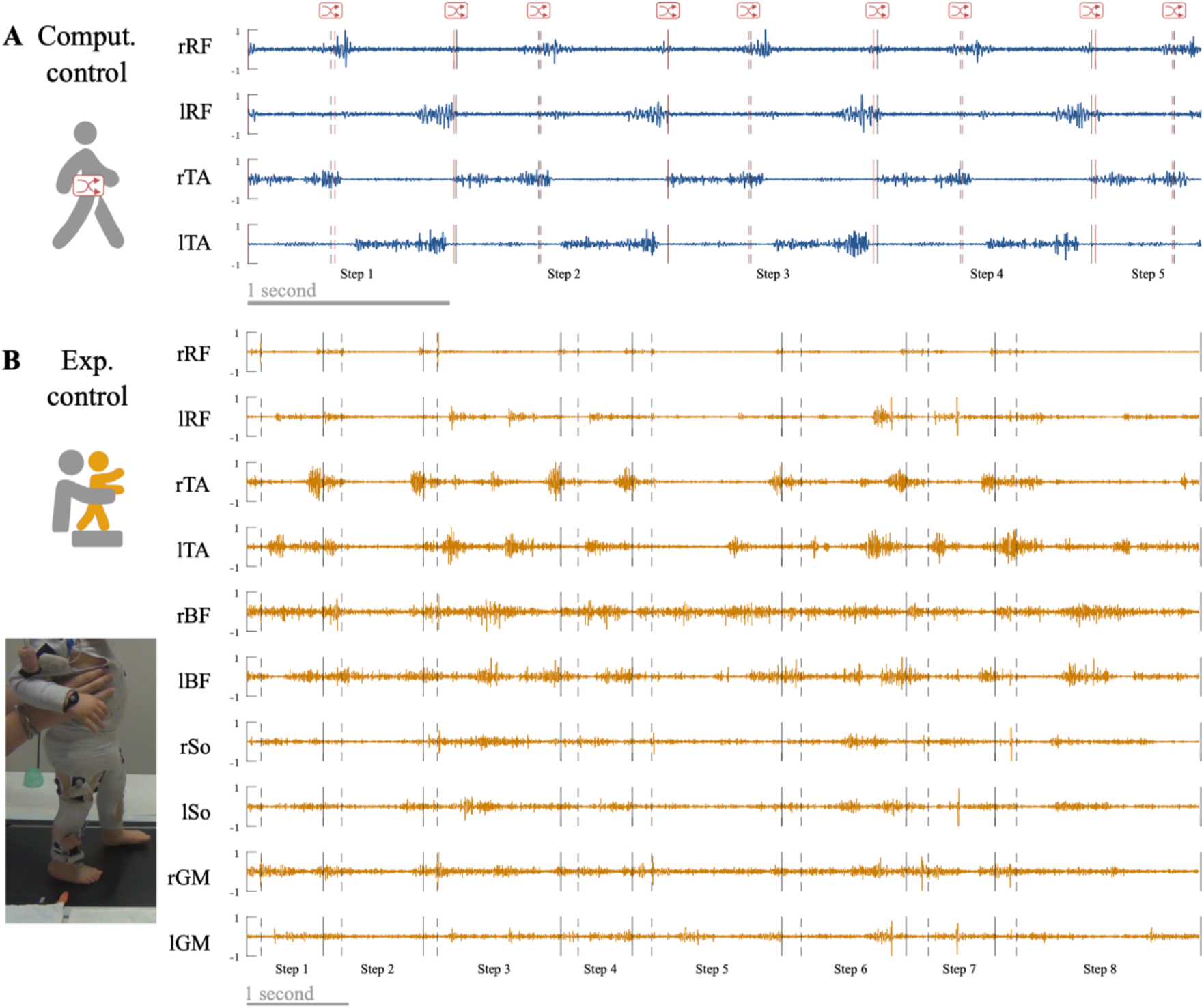
Control conditions. A. Computational control. Raw adult signals are retrieved (this signal is a zoom from Figure 4) and step events are randomly shifted to verify that a difference between adults and toddlers is not due to difficulties to detect step events in toddlers. B. Experimental control. Toddlers were recorded during stepping on a treadmill on the same day that they were recorded during walking in order to verify that variability in toddler was not mainly due to feedback regulations following new balance issues.

The second control condition called “experimental control” verifies the results of toddlers’ data. It was recorded during the same experiment than toddler walking in 10 toddlers. As toddlers are new walkers, their EMG signals could be affected by noise or enhanced EMG variability due to several factors that are not linked with the central command (e.g. walking speed variability, feedback regulations due to postural instability). To address this matter toddlers were recorded during stepping on a treadmill (Figure 3B). During this recording, toddlers were bearing their own weight but moving only their legs while being maintained under the armpits by the experimenter to prevent balance issues. Speed of the belt (and as a result walking speed) was fixed to 17.24 cm/s. We repeated all analyses with those data. As this paper relies on a computational and modeling approach, we cannot draw conclusions regarding the central original of motor modules, however this supplementary analysis controls for the possibility that the results would only originate from feedback regulations of peripheral origin. Individual data for each index in both primary and control conditions are reported in Table *3* (note that no control values exist for toddler 1 and 2 because they did not produce enough stepping cycles).

Each statistical analysis of the paper was first performed between primary conditions (adults vs toddlers values) and was repeated separately for each control condition (i.e. i) replacing adults values with computational control values and comparing it with primary toddlers values, and ii) replacing toddlers values with experimental control values and comparing it with primary adults values, see Table 1 and Table 2).

**Table 1.** Summary of statistical analyses associated with Variant I. Each dataset was modeled with 4 spatial and temporal modules, starting from either averaged or non-averaged EMG data. The resulting VAF (i.e. goodness of fit) was transformed with a Fisher transformation. A mixed ANOVA was performed to characterize the interacting effect of age (adults vs toddlers) and stride-to-stride variability (averaged vs non-averaged data) on modularity. Following this ANOVA, Student t-tests were performed as post-hoc tests to characterize the difference of goodness of fit between adults and toddlers. Comparison between adults and toddlers (first column) is the primary analysis. Other columns report the same analysis with control conditions. Post-hoc p-values were obtained from t-tests and then multiplied by 4 to account for multiple comparisons after the ANOVA (4 post-hoc comparisons).

|  |  | Comparison between adults<br>and toddlers |  | Computational control instead<br>of adult values |  | Experimental control instead<br>of toddlers values |  |
| --- | --- | --- | --- | --- | --- | --- | --- |
|  |  | Non-averaged | Averaged | Non-averaged | Averaged | Non-averaged | Averaged |
| Adults | Mean VAF<br>± SD | 0.74 ± 0.04 | 0.85 ± 0.05 | 0.73 ± 0.04 | 0.85 ± 0.05 | 0.74 ± 0.04 | 0.85 ± 0.05 |
|  | Mean transf.<br>VAF ± SD | 1.30 ± 0.1 | 1.62 ± 0.18 | 1.29 ± 0.09 | 1.64 ± 0.18 | 1.30 ± 0.1 | 1.62 ± 0.18 |
| Toddlers | Mean VAF<br>± SD | 0.57 ± 0.06 | 0.90 ± 0.02 | 0.57 ± 0.06 | 0.90 ± 0.02 | 0.59 ± 0.05 | 0.94 ± 0.02 |
|  | Mean transf.<br>VAF ± SD | 1.00 ± 0.1 | 1.85 ± 0.09 | 1.00 ± 0.1 | 1.85 ± 0.09 | 1.01 ± 0.09 | 2.09 ± 0.15 |
| Mixed ANOVA interacting<br>effect |  | p<.001 |  | p<.001 |  | p<.001 |  |
| Post-hoc t-test p-values<br>(adults vs toddlers) |  | p<.001 | P=.003 | p<.001 | P=.007 | p<.001 | p<.001 |
| Post-hoc t-test p-values<br>(averaged vs non-av.) |  | Adults: p<.001<br>Toddlers: p<.001 |  | Adults: p<.001<br>/ |  | /<br>Toddlers: p<.001 |  |

**Table 2.** Summary of statistical analyses associated with Variant II. We extracted enough modules in each individual to model non-averaged data with a sufficient goodness of fit (i.e. to reach the threshold VAF of 0.75. The mean number of modules for each condition is reported in the table. Then we characterized variability (IRV index) and selectivity of modules activations (IRS index) within the corresponding modular organization. Student t-tests were perform to characterize differences between adults and toddlers. The main analysis is presented on the top line and left columns. To check reliability of results, we repeated the analysis with a fixed number of modules (verification of results with 4 modules, right columns). Every result was verified with control conditions instead of primary ones (see the two last lines).

|  |  | Main analysis (with each individual's number of modules) |  |  | Verification of results with 4 modules |  |
| --- | --- | --- | --- | --- | --- | --- |
|  |  | Number of modules | IRV | IRS | IRV | IRS |
| Mean<br>± SD | Adults | 4.7 ± 0.6 | 4.25 ± 0.93 | 0.61 ± 0.03 | 4.64 ± 0.95 | 0.61 ± 0.02 |
|  | Computational control | 4.7 ± 0.6 | 4.73 ± 1.04 | 0.60 ± 0.03 | 5.16 ± 1.07 | 0.61 ± 0.02 |
|  | Toddlers | 6.8 ± 0.8 | 7.27 ± 1.57 | 0.40 ± 0.04 | 17.28 ± 2.53 | 0.47 ± 0.06 |
|  | Experimental control | 7.1 ± 1.0 | 6.69 ± 2.69 | 0.40 ± 0.04 | 16.77 ± 2.05 | 0.40 ± 0.03 |
| T-test<br>p-values | Adults vs toddlers | p<.001 | p<.001 | p<.001 | p<.001 | p<.001 |
|  | Computational control instead of adult values | p<.001 | p<.001 | p<.001 | p<.001 | p<.001 |
|  | Experimental control instead of toddlers values | p<.001 | p=.01 | p<.001 | p<.001 | p<.001 |

## RESULTS

### Differences in basic kinematics and EMG parameters

Student t-tests on independent samples showed that walking was quite different between the two populations regarding kinematic parameters (Figure 4*C*). As reported by Ivanenko et al. (2013), stride duration was significantly higher in adults than in toddlers walking (p<0.001). However stride duration was significantly higher in experimental control (toddlers stepping) than in adults walking (p<0.001). Variability of stride duration was significantly higher in toddlers than in adults considering either walking (p<0.001) or stepping (i.e. experimental control, p<0.001). Proportion of phases was also significantly different, with adults presenting a shorter proportion of stance phase (63 ± 1.2 %) than toddlers (73.3 ± 4.5 %, p<0.001) with an even longer stance phase in stepping (i.e. experimental control, 79.4 ± 3.6 %, p<0.001).

**Figure 4.**
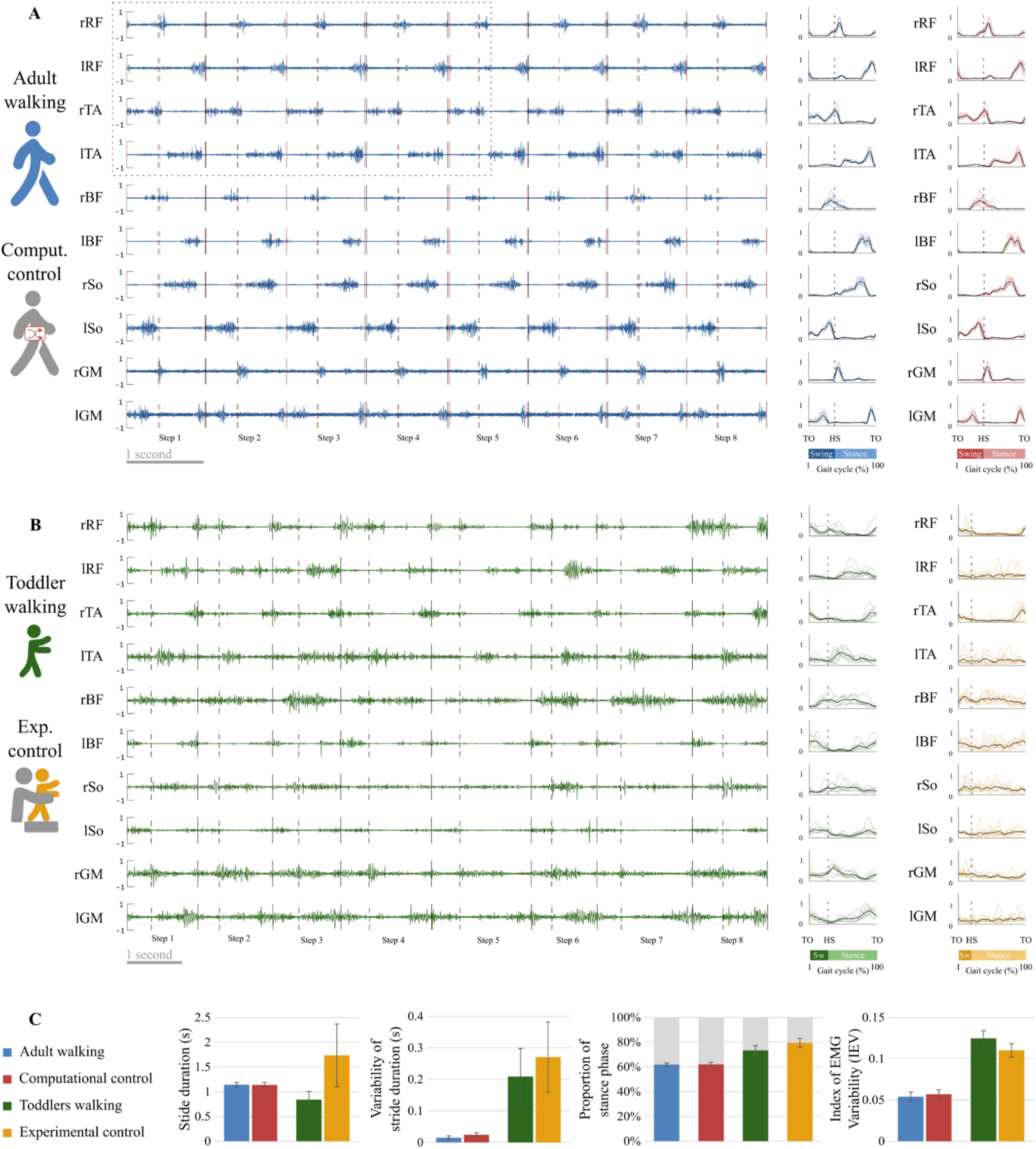
EMG and basic kinematic parameters in adults and toddlers and for control conditions. A. Raw EMG in a representative adult. Preprocessed EMG are depicted at the right of the figure, in blue for this representative adult and in red for the control condition (randomly shifted detection of gait event, see Figure 3A). The black lines represent the average signal of the corresponding muscle in this participant after preprocessing while colored lines represent the signal of the corresponding muscle in each stride for the same participant. B. Raw EMG for a representative toddler. Preprocessed EMG are depicted at the right of the figure, in green for walking and in yellow for the control condition (stepping in the same participant, see Figure 3B). C. Basic kinematic and EMG parameters. From left to right: Stride duration, Variability of stride duration, Proportion of phases across the gait cycle, and Index of EMG Variability. Error bars are standard deviations.

The index of EMG variability (IEV) was significantly higher in toddler than in adults (p<0.001), indicating that pre-processed data were more variable across strides in toddlers. Raw EMG and corresponding pre-processed data are illustrated in Figure 4A and Figure 4B for a representative adult and a representative toddler. Replacing primary adult data by the computational control condition (i.e. adults data with randomly shifted detection of gait events) or toddlers data by the experimental control condition (i.e. toddlers stepping) systematically confirmed this effect (p<0.001, Figure 4*C*, Table *3*).

**Table 3.**
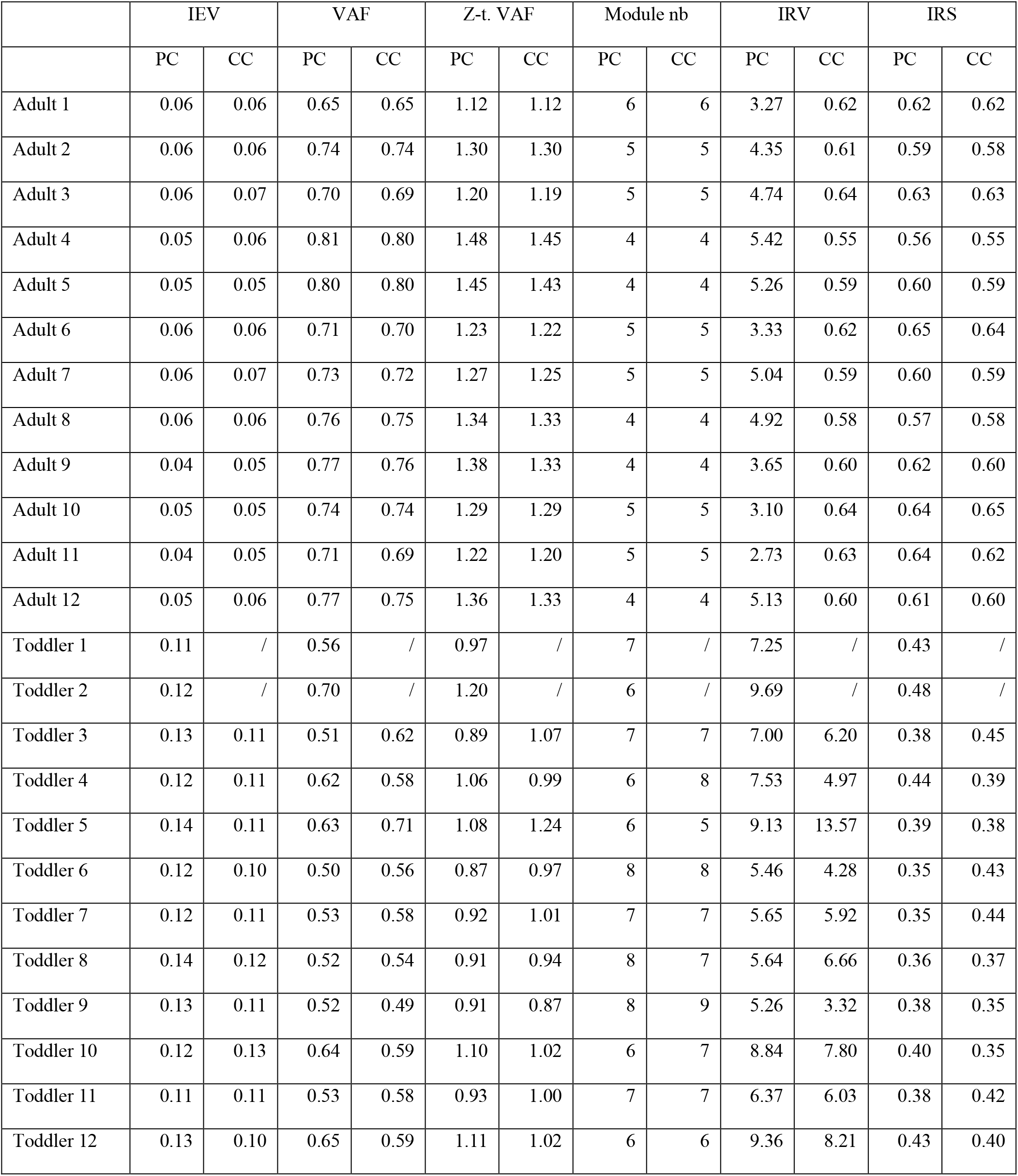
Individual value*s* regarding EMG variability and features of modularity. Each index was computed in Primary Condition (PC) and in Control Condition (CC). CC consists of computational control for adults and of experimental control for toddlers (Figure 3).

### The low-dimensional model of adults modular control does not account for stride-by-stride variability in toddlers

As explained in the method section, we computed the VAF with two approaches: to compare the goodness of fit with a fixed number of modules (variant I), and to compare the modular organization for a fixed goodness of fit (variant II). The results of the first approach are depicted in Figure 5*A*. The mixed ANOVA showed a significant interaction effect between the signal processing (averaged or not) and the population (p<0.001). This result was verified with other methods, extracting only spatial or temporal invariances, or pre-processing data differently (regarding filtering and time-interpolation). Student-t tests were performed as post-hocs and resulting p-values were multiplied by 4 to account for multiple comparisons (4 post-hoc tests). When extracting from an averaged signal across strides, the VAF was higher in toddlers compared to adults (0.90 ± 0.02 vs 0.85 ±0.05). On the contrary, when extracting from a non-averaged signal, the VAF was significantly lower in toddlers compared to adults (0.57 ± 0.06 vs 0.74 ± 0.04, p<0.001). These results show that toddler’s walking EMG signals cannot be decomposed into four spatial and temporal modules with the same goodness of fit as in adults when considering several cycles. Repeating the analysis with both control conditions gave similar values than in primary ones and the ANOVA gave similar results (see Table *1*, Table *3*).

**Figure 5.**
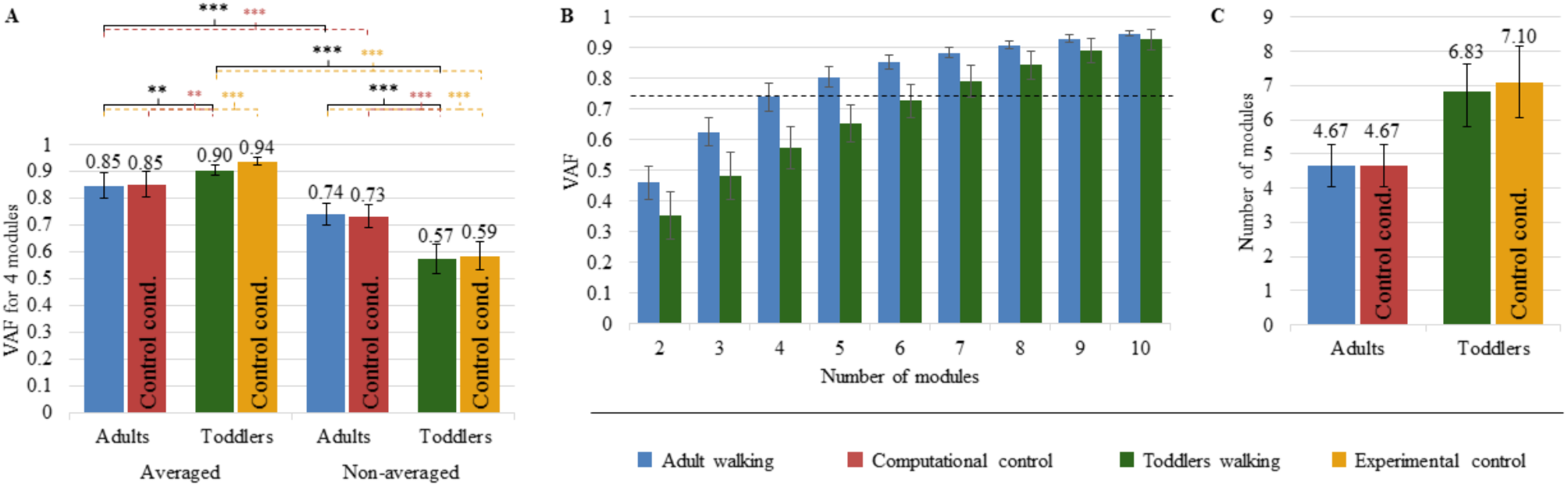
Quality of reconstruction index (VAF) and number of modules. Color code is the same as in Figure 4: Adults and Toddlers primary conditions are respectively depicted in blue and green whereas computational control is depicted in red and experimental control in yellow. A: results from variant I. The plot shows the resulting VAF in adults and toddlers when EMG signals were preprocessed either averaged across strides or not and then factorized into four spatial and temporal modules. Stars show significant differences after post-hoc tests (* p <.05; ** p <.01; *** p<.001). Black stars are for the main analysis and color stars are for repeated analyses with control conditions. B: results from variant II. 2 to 8 modules were extracted from non-averaged data. The plot shows the resulting VAF in adults and toddlers. The dotted line represents the threshold for a good quality of reconstruction (i.e. averaged VAF obtained in adults with four modules). More than 6 modules are necessary on average in toddlers to reach this threshold VAF. C: Number of modules in primary and control conditions.

### Variability and selectivity of toddlers’ modules activation within their higher-dimensional command

The second purpose of our analysis (variant II) was to identify the dimensionality of the modular organization that would allow to reach the same goodness of fit in toddlers than in adults with four modules. The threshold VAF, precisely defined as the averaged VAF obtained in adults by extracting four modules from non-averaged data, was reached differently for each toddler, with on average 6.83 ± 0.80 modules per individual. Applying the same rule for adults resulted in 4.67 ± 0.62 modules per individual which is significantly lower (p<0.001). As such, more modules are needed in toddlers to reach the same quality of reconstruction than in adults walking when considering stride-by-stride variability (Figure 5B, Figure 5C). Nevertheless the number of modules might not be the only interesting feature of a modular organization, as these modules can be activated differently across strides for a high or low number of pairs of spatial and temporal modules. Therefore the indexes of recruitment variability and selectivity (IRV and IRS) were computed following the extraction of the specific number of modules of each individual. Even if extracting from the specific number of modules of each individual only gives a faithful modeling of EMG data, these indexes were also extracted with a fixed number of modules (i.e. following variant I) to verify that the effect was not due to methodological choices.

The IRV was significantly higher in toddlers than in adults, whether it was computed from 4 spatial and temporal modules or from the specific number of modules of each individual (i.e. from variant I or variant II, Figure 6C, p<0.001 in both cases). This indicates that the recruitment of modules occurring on each stride was much more variable in toddlers than in adult, as illustrated in *Figure 6*B by a large spread of activation coefficients across strides. Analyzing data from control conditions instead of primary conditions yielded similar results (Figure 6C, Table 2, Table 3).

**Figure 6.**
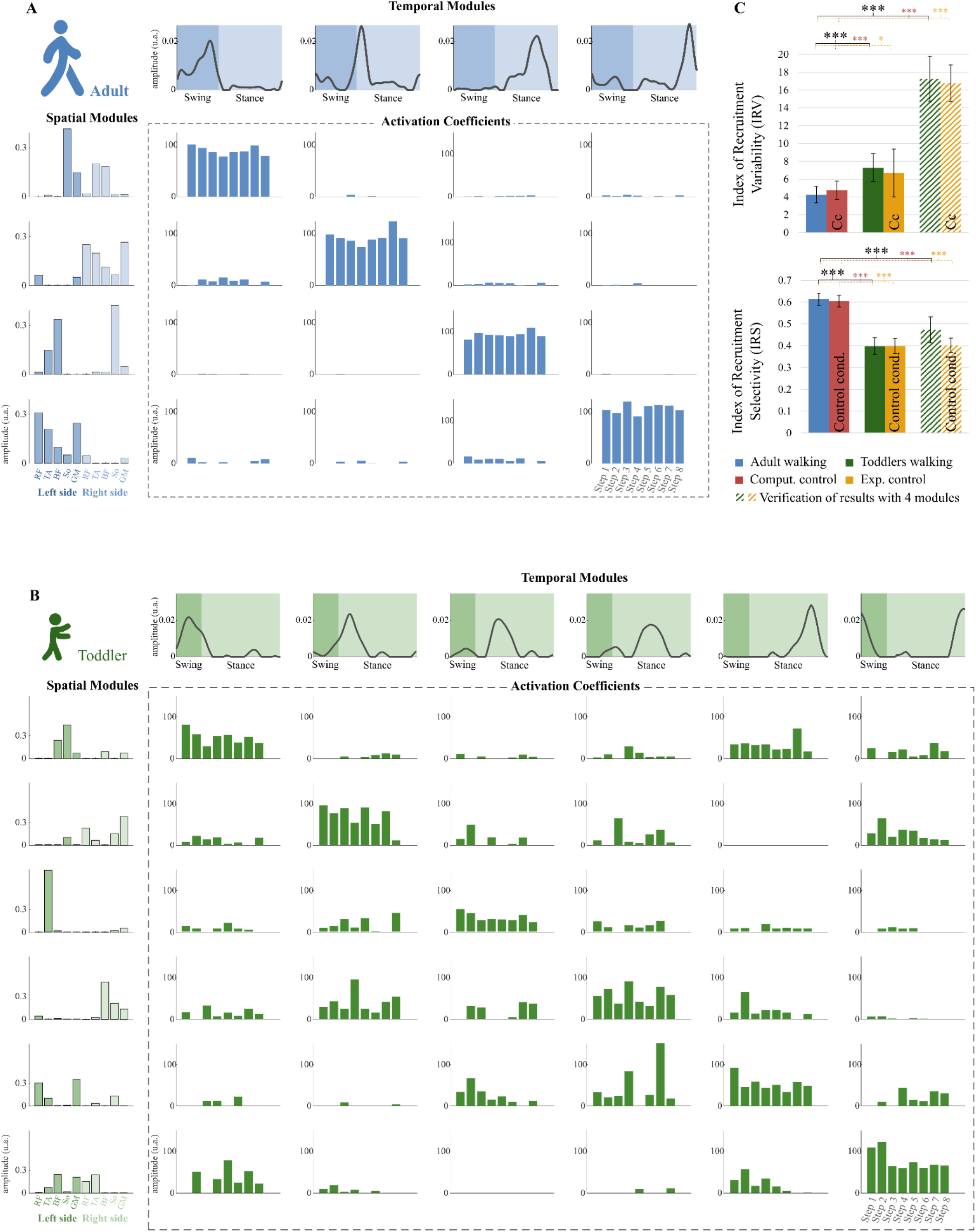
Properties of the modular organization of adults and toddlers. A: Result of the factorization in a representative adult. Adult factorization is depicted for four spatial and temporal modules based on literature. **B:** Result of the factorization in a representative toddler. Toddler factorization is depicted for six spatial and temporal modules, which was the minimum number of modules in this individual to cross the threshold VAF. In each figure, spatial modules are displayed on the left, and muscle weighting are represented in the following order: rectus femoris (RF), tibialis anterior (TA), biceps femoris (BF), soleus (So) and gluteus medius (GM) for the right side then for the left side. Temporal modules are depicted on the top of each figure. They represent 200 time points from the beginning of swing to the end of stance (considering phases of the right lower-limb). Activation coefficients are depicted in the center of each figure. Each bar represents the activation coefficient corresponding to one stride. An activation coefficient represents the concurrent activation of the corresponding pair of spatial and temporal modules during one stride. In each subplot, the y axis represents the amplitude of activation (arbitrary units). **C**. Comparison of the modular organization properties in toddlers and adults. The Index of Recruitment Variability (top) indicates how variables activation coefficients are across the 8 steps in each population and condition. The index of Recruitment Selectivity (bottom) indicates how selectively distributed are those activation coefficients (i.e. exclusively activated with a given spatial or temporal modules or distributed across several ones). Results were verified with a fixed number of modules (stripped bars) to ensure that the effect was not due to methodological choices. Stars show significant differences (* p <.05; ** p <.01; *** p<.001).

The IRS was significantly lower in toddlers, here again from variant I or variant II (Figure 6C, p<0.001 in both cases). The command seems to be more selective in adults than in toddlers. Figure 6 illustrates this difference as it shows that adult modular organization involve four pairs of spatial and temporal modules, each spatial module being activated with one temporal module only (Figure 6A). However, toddler modular organization involves a scattered activation with spatial modules being involves with several temporal modules and vice versa (Figure 6B). Here again, analyzing data from control conditions yielded similar results (Figure 6C, Table *2*, Table 3).

## DISCUSSION

In this paper we aimed to compare the modular organization of toddlers and adults during several strides of walking. Our results showed that the stride-by-stride variability of toddlers’ muscle pattern cannot be simplified in as few dimensions as in adults. From a computational perspective, variability of EMG signals in toddlers seems to involve a high number of computational modules whose degree of activation across steps itself varies, suggesting the plasticity of the motor command. From a neural perspective, these results could indicate the existence of a more complex modularity than in adults, but also the absence of an encoded modularity in toddlers, or the existence of a mixed command associated an adult-like modular organization with other sources of variability. Bellow we discuss these possible interpretations as well as the extent to which modularity could be involved in motor exploration early in development.

Variability appeared as a key feature of the neural control of toddlers at each level of motor organization recorded here: kinematic output (standard error of stride duration), muscular activity (IEV) and activation of modules (IRV). Important variability in muscular activity was already reported in toddlers walking (Chang et al., 2006), stepping (Teulier et al., 2012), or even chewing (Green et al., 1997). Nevertheless it could have been explained by variable activations of a small number of modules that would be equivalent to the one observed in adults, as it seems to be the case in infants (Hinnekens et al. 2022). Adults walking has indeed been described numerous times as controllable with 4 consistently activated modules (Neptune et al., 2009; Clark et al., 2010) with a strong consistency across individuals even when taking into account intra-individual variability (Hinnekens et al., 2020). Here, this 4-modules model fitted well when applied to toddlers’ averaged data, but applying it to toddlers non-averaged data resulted in a significantly lower VAF, be it observed on walking or on the experimental control condition of stepping (with less balance challenge and a fixed speed). Thus the modular activity of toddlers seems to result from a more spread-out and variable activation of a higher number of modules.

On this matter, the first issue to discuss is the potential origin of the variability that is observed in non-averaged data. In general, two types of variability can be distinguished: the one coming from the motor periphery and the one coming from central planning circuits. Peripheral feedback regulations involving reflex loops could play an important role on the muscle activity, thereby limiting our interpretation of the proposed modeling as representing the central command. Nevertheless, we repeated the analysis with a task of stepping on a treadmill with body weight support, which reduced the need for balance-related corrections, and still found a significant stride-by-stride EMG variability in toddlers. Alternatively, centrally generated variability has been widely documented and has been assumed to drive learning-related, purposeful motor exploration (Dhawale et al., 2017). In learning monkeys or songbirds, neurons from different areas of the forebrain seem to generate variability on purpose for motor exploration (Kao et al., 2008; Mandelblat-Cerf et al., 2009). When such motor exploration is prevented by limiting the possible variability of movements, the learning potential at the spinal level is reduced (Ziegler et al., 2010). Similarly in human adults performing a new task, trial-by-trial variability predicts motor learning ability (Wu et al., 2014), and in human infants, the absence of variability can be considered as a sign of motor disability (Hadders-Algra, 2008; Hadders-algra, 2018). Although we cannot fully distinguish the influence of each type of variability on the motor patterns here, there is compelling evidence of its partly central origin and importance for motor exploration which prompts to explore its origins using state-of-the-art models of muscle modularity.

Modeling the motor command at the origin of several strides led to the identification of a higher number of computational modules in toddlers compared to adults. As such, the optimized modular structure that can be found in adults muscle output seems to be shaped and optimized over a long period of time during development. Modules are indeed known to fraction in early life (Dominici et al., 2011; Sylos-labini et al., 2020; Hinnekens et al., 2022), and the results reported here suggest that they could be also merged again at some point between childhood and adulthood. This is indeed coherent with recent studies that identified less motor synergies in walking and/or running in adults than in children (Cheung et al., 2020; Bach et al., 2021). Interestingly, recent data regarding the development of running showed that running started with a small set of computational modules that will first fraction with age and then merge with experience, which respectively corresponds to an increase and a decrease of the number of modules (Cheung et al., 2020). From a computational perspective, the existence of more modules during a temporary state actually makes sense early in development as modularity is associated with a restriction of the possible options for the motor system (Valero-Cuevas, 2009), while variability seems necessary for exploration of the space of possible options (Wu et al., 2014). This temporary state could furnish the possibility to test several muscle associations before choosing the optimized ones. In addition to owning more modules, toddlers seem to be able to flexibly activate those modules across steps, as observed through IRV and IRS indexes (Figure 6). As this leads to an even wider space of possible muscle coordinations, exploration might be particularly boosted during this phase of development. Interestingly this plasticity coincides with an age when infants show a high capacity to shift across locomotor strategies (Ossmy and Adolph, 2020), suggesting an important plasticity of the motor system.

From a neural perspective, a first possible interpretation is the absence of modular control at a temporary point of development. A non-modular control would indeed result in a high number of computational modules after NNMF because the method will always output some modules even if an efficient factorization cannot be established. A second interpretation is that the high dimensionality identified here during several strides corresponds to actual modules that exist within the CNS, and that are fractioned and particularly plastic at this stage of development. In between these two extreme views, walking in toddlers might be the result of a mixed command associating modular inputs that still have to be selected and adequately activated, and non-modular inputs that will be integrated into modules with learning. Mature behaviors are indeed known to rely on shared and task-specific modules, in invertebrates (Jing et al., 2004), frogs (d’Avella and Bizzi, 2005) and humans (Barroso et al., 2014; Nazifi et al., 2017; Hinnekens et al., 2020). Shared modules could be involved when adapting to a new task and facilitate learning before they would be tuned themselves in the longer term (Berger et al., 2013). As such, mature modules might be shaped through practice, as neurons that fire together wire together (Hebb, 1949), and given the particularly important plasticity that exists early in development involving mechanisms such as long-term potentiation that are activity-dependent (de Graaf-Peters and Hadders-algra, 2006; An et al., 2012). Those modules might be tuned during infancy thanks to early spinal plasticity (Vinay et al., 2000; Brumley et al., 2015) which would allow them to remain plastic during toddlerhood. Later in life, mature modules might therefore reflect motor habits originating from coordinations that would have been learned as optimal (de Rugy et al., 2012; Berret et al., 2019).

As stated before, it could be argued that the existence of modules at walking onset could limit motor exploration (Valero-Cuevas, 2009). However, limiting the available space of possible options to some extent might facilitate and guide learning, as described decades ago by Bernstein (1967) and observed nowadays in robotics (Lapeyre et al., 2011) or when using reinforcement learning algorithms, whose success precisely lies on a reduction of the dimensionality of the solution space (Dhawale et al., 2017). Fractioned modules that can still be flexibly activated at walking onset might constitute an ideal compromise of exploration and exploitation. In this vein, the plasticity of motor modules throughout life, associating phases of fractioning and merging of modules (Cheung et al., 2020; Sylos-labini et al., 2020; Hinnekens et al., 2022) might follow the development of other sources of constraints. For example in the womb, the limited space constrains possibilities for the kicking behavior (Piek, 2002) which leads to the emergence of a specific kicking pattern available at birth (Musselman and Yang, 2008; Robinson et al., 2008). In this context, modules could be used to store temporary solutions, creating new constraints at time points when external constraints are diminished. As such, development of modularity is likely to be a dynamic mechanism alternating phases of module shaping and phases of exploration within the resulting restricted space, as “in a modular controller, learning is partitioned into two processes: learning the modules and learning the parameters of the modules’ combination rules” (d’Avella and Pai, 2010). The modular system should continue to be tuned until the end of growth, as mature modules need to integrate biomechanical properties of the musculoskeletal system (Bizzi et al., 1991; Bizzi and Cheung, 2013) as well as each individual’s specificities (Torres-Oviedo and Ting, 2010; Bizzi and Cheung, 2013). Although descriptive and based on EMGs, our study suggests that the motor system is quite plastic around toddlerhood, coherently with current recommendations for the implementation of early therapies (Ulrich, 2010). Future studies need to better identify if critical periods of module acquisition exist during motor development.

## COMPETING INTERESTS

The authors declare no competing interests.

## DATA AND CODE AVAILABILITY

Individual data supporting the findings of this study are available in Table 3. Additional data related to this paper may be requested from the authors. Custom code used in this study is available at: http://hebergement.universite-paris-saclay.fr/berret/software/sNM3F.zip

## ACKNOWLEDGMENTS

We thank Prof. François Goffinet, head of the Port-Royal maternity in Paris, for encouraging this study. We also thank the Région Ile-de-France for their participation in the initial set-up of the babylab. Finally, we warmly thank all infants and parents who participated in the study.

## Notes

### Competing Interest Statement

The authors have declared no competing interest.

